# Heat Acclimation in Mice Requires Preoptic BDNF Neurons and Postsynaptic Potentiation

**DOI:** 10.1101/2024.11.13.623142

**Authors:** Baoting Chen, Cuicui Gao, Changhao Liu, Tongtong Guo, Junwei Hu, Jialiang Xue, Kangmin Tang, Yuelai Chen, Tian Yu, Qiwei Shen, Hongbin Sun, Wen Z. Yang, WeiL. Shen

**Author notes:** Corresponding authors: H.S., W.Z.Y; W.L.S; (https://slst.shanghaitech.edu.cn/sw_en/main.htm). These authors contributed equally (Baoting Chen, Cuicui Gao).

## Abstract

Heat acclimation (HA) is a key adaptive response in mammals to repeated heat exposure, essential for fitness and survival^1-3^. HA improves cardiovascular function, thermal comfort, and exercise capacity^4, 5^. However, the lack of a genetically tractable model has hindered understanding of the molecular and neural mechanisms underlying HA. Here, we show that 10 days of daily 38°C exposure lowers core body temperature (T_core_) and reduces anxiety during subsequent heat exposures in mice. HA increases brain-derived neurotrophic factor (BDNF) expression in the medial preoptic area (MPO). BDNF-expressing MPO neurons (MPO^BDNF^) show increased intrinsic heat sensitivity after HA. These neurons orchestrate downstream targets in the dorsomedial hypothalamus (DMH) and rostral raphe pallidus (rRPa) to mediate HA effects. BDNF, acting through its receptor tropomyosin-related kinase B (TrkB) in the DMH, facilitates the anxiolytic effect of HA by enhancing excitatory synaptic connections between MPO^BDNF^ and DMH neurons. This study provides new insights into HA mechanisms, setting the stage for future research on heat stress reduction and exercise optimization.

## Result

Without HA, heat exposure (38°C) increased anxiety but did not impair novel object recognition in male *C57* mice, as tested in the open field test (OFT) and Y-maze (Supplementary Information, Fig. S1a, b). To establish an effective HA protocol, we evaluated three different protocols. Neither daily exposures to 35°C for 3 hours over 14 days nor daily exposures to 35-38°C for 3 hours over 20 days impacted T_core_ during subsequent 38°C exposures (Supplementary Information, Fig. S1c, d). However, consecutive 10-day exposures to 38°C for 2 hours significantly improved heat tolerance, as marked by a moderate rise in T_core_ during subsequent 38°C exposures (Supplementary Information, Fig. S1e). Thus, we established an effective HA protocol. Under this protocol, food intake decreased while water intake significantly increased during the 2-hour heat exposure. Over the full 10-day acclimation period, water consumption increased significantly, while no changes were observed in body weight or food intake (Supplementary Information, Fig. S1f-l).

After HA, we re-exposed the mice to 38-40°C to systematically assess the adaptive effects (Fig. 1a). HA mice displayed extended T_core_ tolerance in the heat tolerance test (HTT) at 40°C, lasting three times longer than in the non-HA group, as measured by the latency for T_core_ to reach 41.5°C (a critical thermal limit^6, 7^) (Fig. 1b). Also, HA mice exhibited increased open arm exploration in the elevated plus maze (EPM) (Fig. 1c) and central exploration during OFT (Supplementary Information, Fig. S2a, b), indicative of reduced anxiety. However, HA did not influence thermal preference or nociceptive heat perception (Supplementary Information, Fig. S2c, d). While physical activity remained constant during heat exposure at 38°C, energy expenditure (EE) reduced (Supplementary Information, Fig. S2e, f). This reduction in EE aligned with decreased mRNA levels of uncoupling protein 1 (UCP1) in brown adipose tissue (Supplementary Information, Fig. S2g). HA did not affect the time spent in or the number of entries into the closed arms in the Y-maze (Supplementary Information, Fig. S2h). HA also enhanced thermoregulatory behaviors, characterized by longer grooming time (Supplementary Information, Fig. S2i) and faster, longer body postural extension during heat exposures (Supplementary Information, Fig. S2j). Similarly, HA training in female mice also enhanced T_core_ tolerance and reduced heat-induced anxiety (Supplementary Information, Fig. S3a-c). Thus, HA significantly improves heat tolerance and alleviates anxiety by reducing thermogenesis and bolstering thermoregulatory capacity, without affecting cognitive and pain perception functions.

**Fig. 1:**
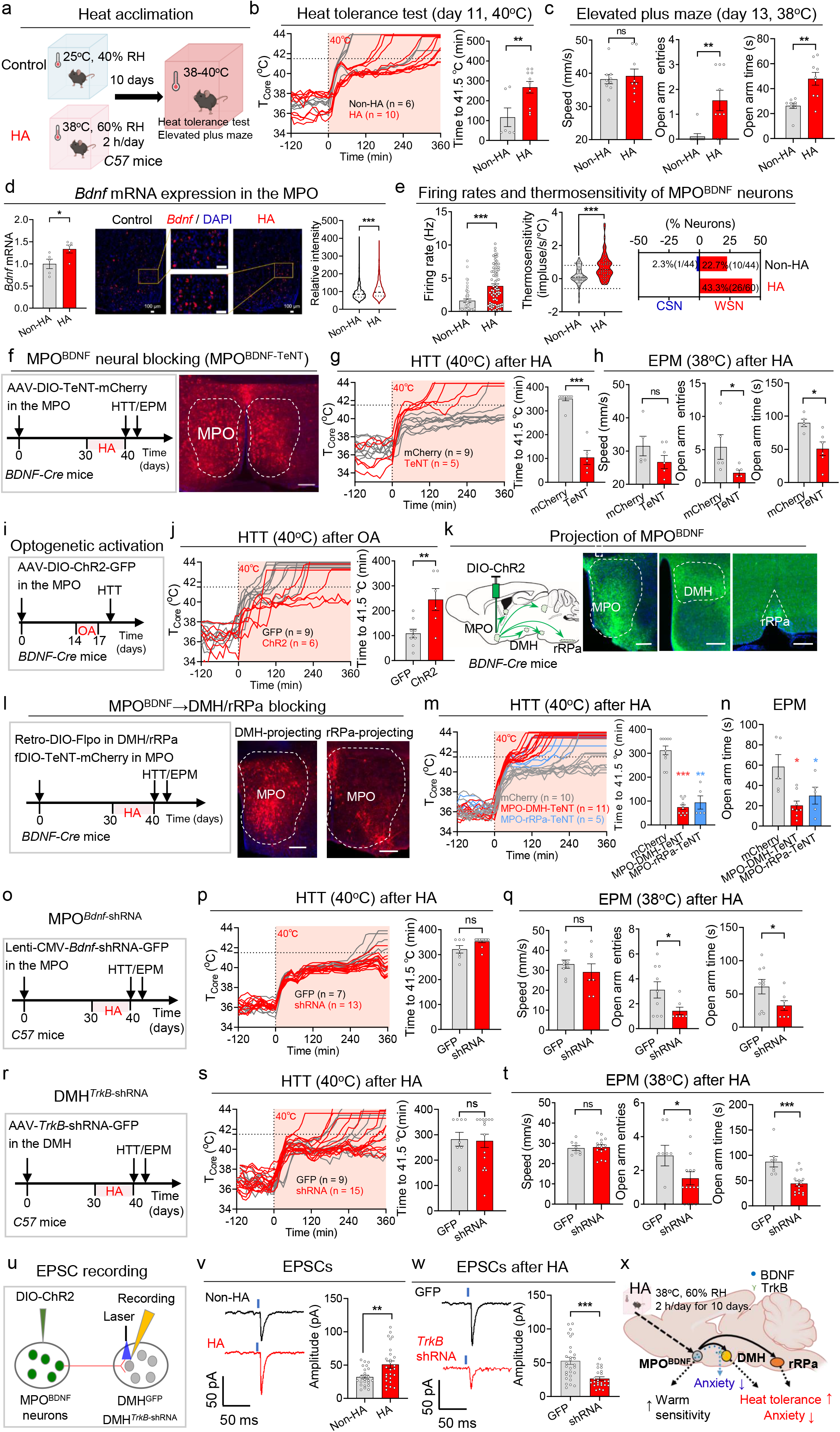
BDNF in the MPO is required for heat acclimation. **a** The experimental procedure for HA. The *C57* mice were subjected to controlled conditions of 25oC and 40% relative humidity (RH) in an incubator. The HA group underwent a 10-day training program consisting of daily 2-hour sessions under conditions of 38oC and 60% RH. HTT and EPM were conducted after HA. **b** Body temperature (T_core_) changes in exposure to ambient temperature of 40oC in the HTT from individual mice. Some mice would die after prolonged exposure to 40°C, and their T_core_ after death were replaced with the highest T_core_ to produce a flattened trace. The time taken for T_core_ to reach 41.5 oC (the critical T_core_) was quantified in the right. **c** EPM under heat exposure (38oC). The traveling speed, entries into open arm and time spent in open arm were quantified. **d** The expression of *Bdnf* mRNA in the MPO in HA mice in response to a 2-hour heat exposure (38oC), determined by qPCR (left) and *RNAscope* (middle). The *Bdnf* mRNA expression of each neuron is quantified in the right. Scale bar, 100 μm. **e** The basal firing rates at 36oC, the thermosensitivity of BDNF_+_ neurons in the MPO (termed as MPO_BDNF_) and the percentage of warm-sensitive neurons (WSN) or cold-sensitive neurons (CSN) within the recorded BDNF_+_ neurons. **f** Protocol for HA training after blocking MPO_BDNF_ neurons with TeNT and the representative expression of AAV-DIO-TeNT-mCherry. AAV-DIO-TeNT was injected in the MPO of the *BDNF-IRES-Cre* mice (termed as *MPO*_*BDNF-TeNT*_, TeNT group). The AAV-DIO-mCherry (mCherry group) was used as the control. Scale bar, 200 μm. **g, h** HTT (**g**) and EPM (**h**) after HA training in *MPO*_*BDNF-TeNT*_ mice. **i** Experimental design to test the effects of optogenetic activation (OA) of MPO_BDNF_ neurons. AAV-DIO-ChR2-GFP was injected in the MPO of the *BDNF-IRES-Cre* mice (termed as *MPO*_*BDNF-ChR2*_, ChR2 group). The AAV-DIO-GFP (GFP group) was used as the control. **j** HTT of the *MPO*_*BDNF-ChR2*_ mice after OA. **k** Axonal GFP expression of MPO_BDNF-ChR2-GFP_ neurons in the DMH and RPa. Scale bar, 200 μm. **l** Protocol for HA training after blocking DMH/rRPa-projecting MPO_BDNF_ neurons and representative expression of AAV-fDIO-TeNT-mCherry. Retro-DIO-Flpo was injected in the DMH/rRPa and AAV-fDIO-TeNT-mCherry was injected in the MPO of the *BDNF-IRES-Cre* mice (MPO-DMH/rRPa TeNT group). The AAV-fDIO-mCherry (mCherry group) was used as the control. Scale bar, 200 μm. **m, n** HTT (**m**) and EPM (**n**) after HA training in MPO-DMH/rRPa TeNT group. **o** Design to knockdown *Bdnf* in MPO by injecting Lenti-CMV-*Bdnf*-shRNA into *C57* mice (termed as *MPO*_*BDNF-shRNA*_, shRNA group). The scrambled shRNA group was used as the control. **p, q** HTT (**p**) and EPM (**q**) after HA training in *MPO*_*Bdnf-shRNA*_ mice. **r** Design to knockdown *TrkB* in DMH neurons by injecting AAV-hSyn-Cre & AAV-DIO-shRNA(*TrkB*)-GFP into *C57* mice (termed as *DMH*_*TrkB-shRNA*_, shRNA group). The mice injected AAV-hSyn-Cre & AAV-DIO-shRNA(*NC*)-GFP (GFP group) were used as the control. **s, t** HTT (**s**) and EPM (**t**) after HA training in *DMH*_*TrkB-shRNA*_ mice. **u** Experimental design to record postsynaptic currents in DMH neurons innervated by MPO_BDNF_ neurons using whole-cell patch configurations. **v** Representative traces and amplitudes of EPSCs recorded in DMH neurons following light stimulation (473 nm, 5 ms) of MPO_BDNF_ afferents. The *MPO*_*BDNF-ChR2*_ mice without HA training were as the control group. **w** Representative traces and amplitudes of EPSCs recorded in DMH neurons following light stimulation. AAV-hSyn-*TrkB*-shRNA were injected in the DMH of *MPO*_*BDNF-ChR2*_ mice. The scrambled shRNA group was used as the control. **x** Summary of the role of BDNF in HA. HA, heat acclimation; HTT, heat tolerance test; EPM, elevated plus maze; BDNF, brain-derived neurotrophic factor; MPO, medial preoptic area; DMH, dorsomedial hypothalamus; rRPa, rostral raphe pallidus; TrkB, tropomyosin-related kinase B; EPSC, excitatory postsynaptic current. All data are shown as mean ± SEM and analyzed by *t*-test. *p < 0.05, **p < 0.01, ***p < 0.001 vs corresponding control group. ns, not significant.

Building on previous findings that BDNF neurons in the POA are crucial for heat defense and that acute heat exposure significantly increases POA BDNF expression^8, 9^, we hypothesized that BDNF plays a role in HA. In line with this, HA mice showed elevated *Bdnf* mRNA expression after subsequent 38°C exposure (Fig. 1d). Considering the importance of intrinsic thermosensitivity in heat defense^10, 11^ and its proposed role in HA^12, 13^, we first recorded the basal firing rates at 36°C and found that HA notably increased the firing rates of MPO^BDNF^ neurons. We further found that the thermosensitivity of MPO^BDNF^ neurons increased significantly after HA, resulting in a rise in the ratio of warm-sensitive neurons (WSNs, defined as neurons with thermosensitivity > 0.8 imp/s/°C). The ratio increased from 22.7% to 43.3% (Fig. 1e), exceeding the overall increase in MPO neurons (from 27.3% to 36.7%; Supplementary Information, Fig. S4a). Taken together, these results highlight that HA increases MPO BDNF expression, and intrinsic thermosensitivity of MPO^BDNF^ neurons.

To determine the roles of MPO^BDNF^ neurons in HA, we inhibited them using tetanus neurotoxin (TeNT) (Fig. 1f). Consistent with their reported role in heat defense^9^, this inhibition impaired basal heat defense function, leading to a rise in T_core_ at 38°C during the first day of HA training (Supplementary Information, Fig. S4b, c). Notably, this inhibition nearly abolished the HA effect on T_core_ tolerance, with T_core_ rapidly increasing to 41.5°C during the HTT (Fig. 1g). Concurrently, the anxiety-reducing effect of HA was also negated (Fig. 1h; Supplementary Information, Fig. S4d). Thus, MPO^BDNF^ neurons are required for the HA effects on T_core_ and anxiety.

To ascertain whether MPO^BDNF^ neural activation could induce an HA effect without HA training, we employed optogenetic activation using ChR2 (Fig. 1i). Intriguingly, optostimulation at 1 Hz over three consecutive days elicited a pronounced HA effect, evidenced by a two-fold increase in the time taken to reach 41.5°C in the HTT compared with GFP controls (Fig. 1j). To identify downstream targets of MPO BDNF neurons, we considered the DMH and rRPa (Fig. 1k), as suggested in the thermoregulatory circuitry^14^. We then employed a projection-specific strategy to block synaptic transmission (Fig. 1l). As expected, blocking either DMH- or rRPa-projecting MPO^BDNF^ neurons had a small but significant effect on basal heat defense function during 38°C exposures (Supplementary Information, Fig. S4e, f). Notably, blocking either of these two pathways nearly abolished the HA effect on T_core_ tolerance and anxiety alleviation during heat exposures (Fig. 1m, n; Supplementary Information, Fig. S4g, h). These data collectively demonstrate that the MPO^BDNF^→DMH/rRPa neurocircuitry is essential for the HA effect.

To determine the role of BDNF in HA, we knocked down its expression by injecting Lenti-*Bdnf*-shRNA virus into the MPO of *C57* mice (Fig. 1o). BDNF knockdown in the MPO did not alter the basal heat defense function (Supplementary Information, Fig. S4i-k). Surprisingly, BDNF knockdown had no effect on T_core_ tolerance during HTT (Fig. 1p). In contrast, the knockdown compromised the anxiolytic effects of HA, as indicated by reduced open arm exploration in the EPM (Fig. 1q) and reduced central exploration in the OFT (Supplementary Information, Fig. S4l).

Therefore, BDNF in the MPO is essential for the anxiolytic effects of HA.

To determine whether the BDNF receptor TrkB is involved in HA, we knocked it down in various regions axonally projected by MPO^BDNF^ neurons, including the DMH, rRPa, paraventricular thalamus (PVT), periaqueductal grey (PAG), and mamillary peduncle (mp) (Fig 1r; Supplementary Information, Fig. S5a, b). Similar to BDNF knockdown in the MPO, TrkB knockdown in the DMH did not affect basal heat defense when exposed to 38°C (Supplementary Information, Fig. S5c-e). Additionally, it had no effect on T_core_ tolerance during HTT (Fig. 1s). However, it significantly reduced the anxiolytic effects of HA, as evidenced by decreased open arm exploration in the EPM (Fig. 1t) and reduced central exploration in the OFT (Supplementary Information, Fig. S5f). Beyond the DMH, knockdown in none of other regions—rRPa, PVT, PAG, or mp—affected HA’s impact on T_core_ or anxiety (Supplementary Information, Fig. S5g-j). Also, the simultaneous knockdown of TrkB in the DMH and rRPa did not affect T_core_ tolerance (Supplementary Information, Fig. S5k). In conclusion, TrkB in the DMH is selectively required for the anxiolytic effects of HA.

Synaptic plasticity is postulated as a pivotal mechanism underlying HA^12^. Given that BDNF is known to modulate synaptic plasticity^15^, we suspected that BDNF-dependent synaptic remodeling within the BDNF circuitry could be instrumental for HA. Previous studies documented that MPO^BDNF^ neurons consist of mixed glutamatergic and GABAergic subpopulations, with proportions of 60% and 33%, respectively^8, 9^. We confirmed both connections by recording excitatory and inhibitory postsynaptic currents (EPSCs/IPSCs) in DMH neurons, evoked by optogenetic stimulation of MPO^BDNF^ terminals in the DMH (Fig. 1u; Supplementary information, Fig. S6a). Our data showed 64-67% of neurons had light-evoked EPSCs, 62% had light-evoked IPSCs, and 26-29% exhibited both EPSCs and IPSCs in either HA or control groups (Supplementary information, Fig. S6b). However, the amplitude of light-evoked EPSCs increased (Fig. 1v), while that of IPSCs decreased in HA mice (Supplementary information, Fig. S6c), suggesting potentiation of excitatory and depression of inhibitory connections in the MPO^BDNF^→DMH pathway.

To discern whether this potentiation was pre- or post-synaptic, we analyzed the paired-pulse ratio (PPR) of two light-evoked EPSCs within a 50-ms interval and characterized properties of miniature EPSCs/IPSCs. There was no difference in PPR or mEPSCs frequency between groups, indicating no change in presynaptic glutamate release probability (Supplementary information, Fig. S6d, e). However, there was a marked increase in the amplitude of mEPSCs (Supplementary information, Fig. S6e), pointing to an increase in postsynaptic ion conductance. In contrast, both the amplitude and frequency of mIPSCs remained unchanged (Supplementary information, Fig. S6f). Further quantal EPSCs recordings with strontium (Sr)^16^ revealed that HA mice exhibited increased amplitude but unchanged frequency of quantal EPSCs compared to controls (Supplementary information, Fig. S6g). Knocking down TrkB in DMH neurons abolished HA-mediated EPSC potentiation (Fig. 1w; Supplementary information, Fig. S6h) but did not affect IPSC amplitude (Supplementary information, Fig. S6i). Taken together, we conclude that the MPO^BDNF^→DMH pathway undergoes TrkB-dependent potentiation of excitatory transmissions primarily via increased postsynaptic ion conductance.

In summary, we establish a simple and easily replicable heat acclimation (HA) protocol in mice and explore HA’s profound effects on thermoregulation and anxiety alleviation. These effects are primarily mediated by the MPO^BDNF^→DMH/rRPa thermoregulatory pathways (Fig. 1x). Within these pathways, HA enhances BDNF expression in the MPO and increases neuronal intrinsic thermosensitivity. These BDNF molecules interact with TrkB in the DMH, potentiating excitatory synaptic connectivity and alleviating anxiety. Additionally, it is worth noting that BDNF levels in human serum also increase following heat exposure^17^, suggesting a potentially conserved mechanism between mice and humans. Thus, our research not only deepens our understanding of physiological adaptations to heat but also informs strategies to improve thermal performance.

This article has been accepted by *Cell Research*.

## Acknowledgments

This study was funded by National Nature Science Foundation of China (9235730017, 32330042, and 32425028 to W. Shen; 32100825 to W. Yang), Shanghai Sailing Program (21YF1429800 to W. Yang), Shanghai Science and Technology Committee of Shanghai City (19140903800 and 21XD1422700 to W. Shen), and Shanghai Frontiers Science Center for Biomacromolecules and Precision Medicine at ShanghaiTech University. We thank the staff members of the Animal Facility at ShanghaiTech University for providing technical support.

## Author contributions

B.C., and C.G performed most of the experiments; C.L performed electrophysiology; J.X., and J.H. helped data analysis; Y.C., K.T., T.Y., Q.S., H.S., W.Z.Y. and W.L.S. designed the experiments; H.S., B.C., W.Z.Y., and W.L.S. generated the figures. W.Z.Y., H.S., and W.L.S. wrote the manuscript.

## Data and materials availability

All data presented in the main text or supplementary materials are available from the corresponding author upon reasonable request.

## Reference

1 Kenny GP, Wilson TE, Flouris AD, Fujii N. Heat exhaustion. Handb Clin Neurol 2018; 157:505–529.

2 Qiu M, Zhou Y, Ye N et al. Improved Heat Tolerance in Heat-acclimated Mice: The Probable Role of the PD-L1 Pathway. bioRxiv 2022:2022.2002.2020.481185.

3 Habibi P, Razmjouei J, Moradi A, Mahdavi F, Fallah-Aliabadi S, Heydari A. Climate change and heat stress resilient outdoor workers: findings from systematic literature review. BMC Public Health 2024; 24:1711.

4 Horowitz M. Heat acclimation, epigenetics, and cytoprotection memory. Comprehensive Physiology 2014; 4:199–230.

5 Taylor NA. Human heat adaptation. Comprehensive Physiology 2014; 4:325–365.

6 Wilkinson DA, Burholt DR, Shrivastava PN. Hypothermia following whole-body heating of mice: effect of heating time and temperature. Int J Hyperthermia 1988; 4:171–182.

7 Koh YH. Heat Stroke with Status Epilepticus Secondary to Posterior Reversible Encephalopathy Syndrome (PRES). Case Rep Crit Care 2018; 2018:3597474.

8 Zhao ZD, Yang WZ, Gao CC et al. A hypothalamic circuit that controls body temperature. Proceedings of the National Academy of Sciences of the United States of America 2017; 114:2042–2047.

9 Tan CL, Cooke EK, Leib DE et al. Warm-Sensitive Neurons that Control Body Temperature. Cell 2016; 167:47-+.

10 Zhou Q, Fu X, Xu J et al. Hypothalamic warm-sensitive neurons require TRPC4 channel for detecting internal warmth and regulating body temperature in mice. Neuron 2023; 111:387–404 e388.

11 Boulant JA, Dean JB. Temperature receptors in the central nervous system. Annual review of physiology 1986; 48:639–654.

12 Armstrong LE, Stoppani J. Central nervous system control of heat acclimation adaptations: an emerging paradigm. Rev Neurosci 2002; 13:271–285.

13 Horowitz M. Heat acclimation: phenotypic plasticity and cues to the underlying molecular mechanisms. J Therm Biol 2001; 26:357–363.

14 Morrison SF, Nakamura K. Central neural pathways for thermoregulation. Frontiers in bioscience 2011; 16:74–104.

15 Wang CS, Kavalali ET, Monteggia LM. BDNF signaling in context: From synaptic regulation to psychiatric disorders. Cell 2022; 185:62–76.

16 Grzelka K, Wilhelms H, Dodt S et al. A synaptic amplifier of hunger for regaining body weight in the hypothalamus. Cell metabolism 2023.

17 Kojima D, Nakamura T, Banno M et al. Head-out immersion in hot water increases serum BDNF in healthy males. Int J Hyperthermia 2018; 34:834–839.

