## Supplemental Fig1-6 for "Heat Acclimation in Mice Requires Preoptic BDNF Neurons and Postsynaptic Potentiation"

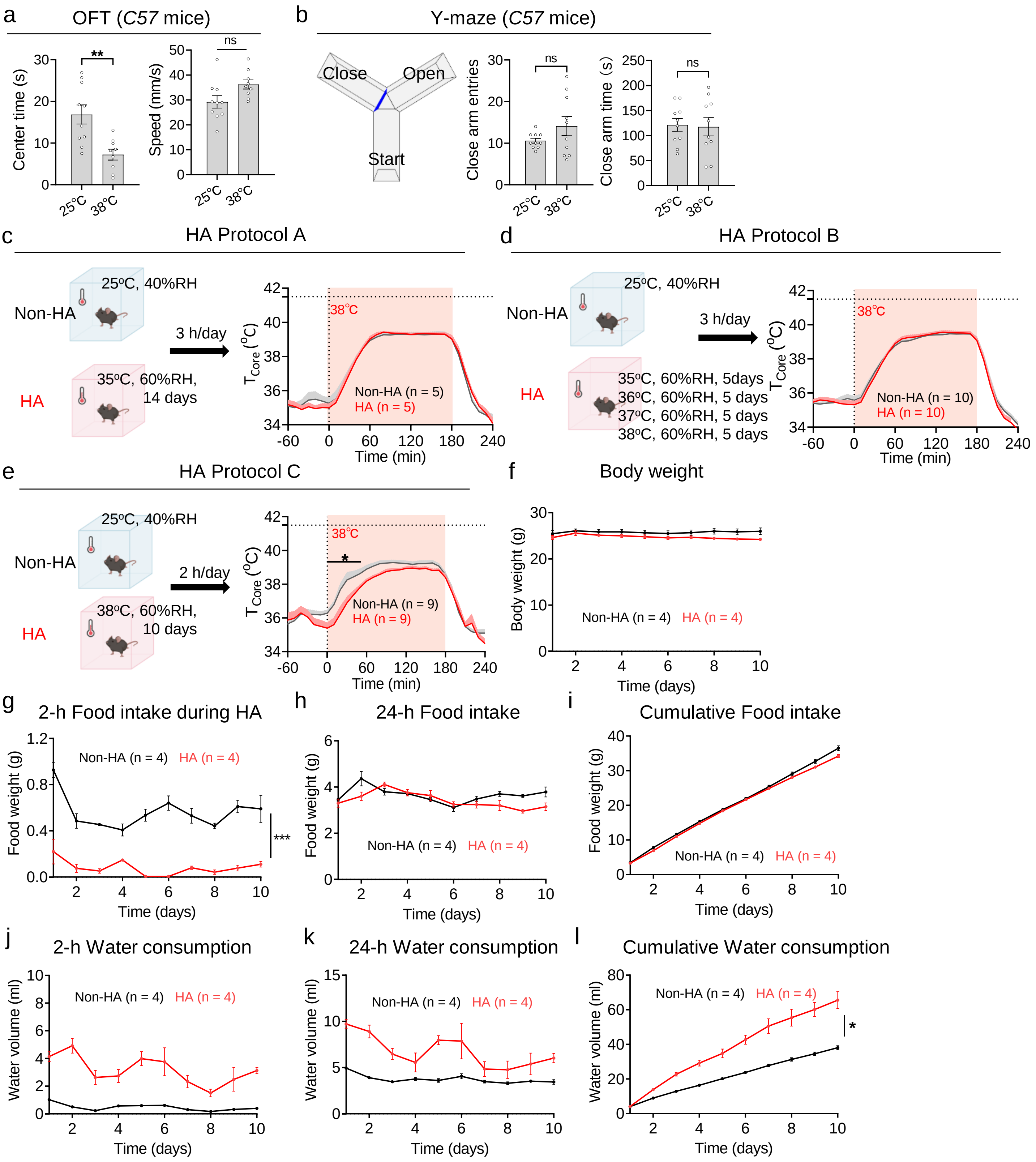


**Fig. S1. Behavioral changes in *C57* mice after heat exposure.**

**a, b** Open field test (OFT) (**a**) and Y-maze (**b**) performed under 25ºC and 38ºC in *C57* mice. In OFT, the time spent in the center zone and the traveling speed in 5-min were quantified. In Y-maze, the entries into close arm and time spent in close arm in 5-min were quantified.

**c-e** Different protocols for HA trainings as indicated in figures. All the HA mice were under conditions of 60% RH. $T_{core}$ changes in exposure to 38ºC after HA training was used to assess the outcomes of the training.

**f** Changes in body weight during HA training, using protocol C.

**g-i** During HA training, the amount of food consumed was documented for different time periods: 2-hour(**g**), 24-hour (**h**), and 10-day (**i**).

**j-l** During HA training, the amount of water consumed was documented for different time periods: 2-hour(**j**), 24-hour (**k**), and 10-day (**l**).

All data are shown as mean ± SEM and analyzed by *t*-test (**a, b**) and two-way RM ANOVA (**c-l**). *p < 0.05, **p < 0.01, ***p < 0.001 vs corresponding control group. ns, not significant.

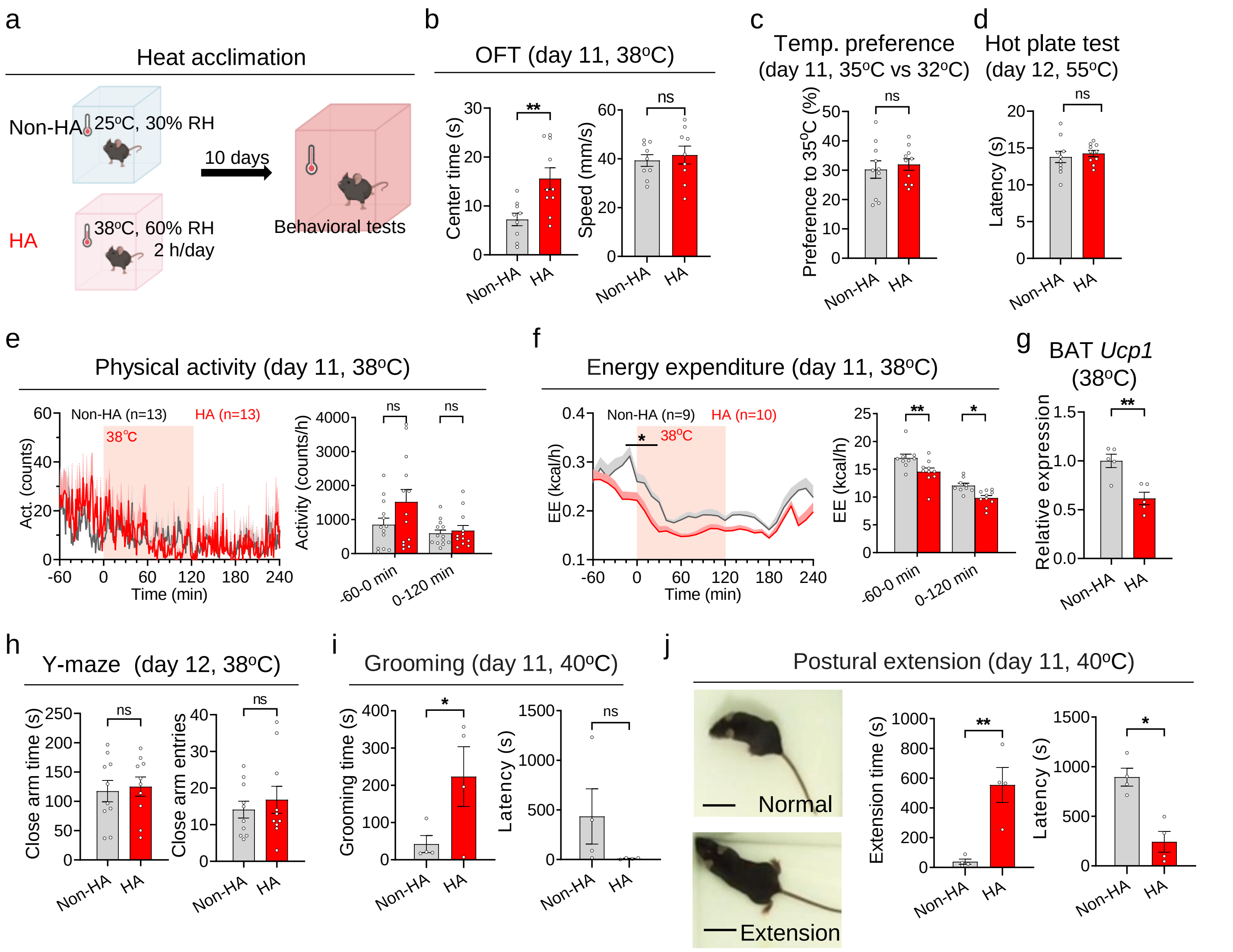


**Fig. S2. Phenotypes changes in HA mice.**

**a** The experimental procedure for HA.

**b** OFT under heat stress (38ºC).

**c** Temperature preference test after HA training. The time ratio mice spent in 35ºC within 30min was counted.

**d** Hot plate tests (60ºC) after HA training. Latency for paw withdrawal was recorded.

**e, f** Changes in physical activity (**e**) and energy expenditure (**f**) after HA training in response to heat exposure (38ºC).

**g** The *Ucp1* mRNA expression in BAT in HA and Non-HA mice in response to a 2-hour heat exposure (38ºC).

**h** Y-maze after HA training under heat stress (38ºC).

**i** Grooming behavior after HA training under heat stress (40ºC). The time and latency of onset were quantified.

**j** Postural extension after HA training under heat stress (40ºC). The representative images (left), the time (middle), and latency of onset (right) were shown. Scale bar, 5 cm.

All data are shown as mean ± SEM and are analyzed by *t*-test (**b-d, g-j**) or two-way RM ANOVA (**e-f**). *p < 0.05, **p < 0.01, ***p < 0.001 vs corresponding control group. ns, not significant.

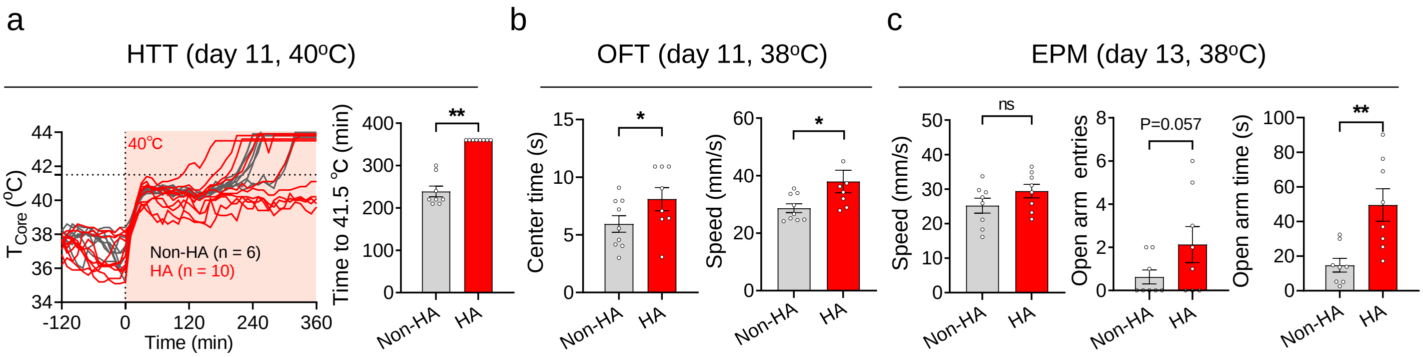


**Fig. S3. Behavioral changes in *C57* female mice after HA training.**

**a** HTT after HA training in *C57* female mice.

**b, c** OFT (**b**) and EPM (**c**) performed under heat exposure (38ºC) in *C57* female mice.

All data are shown as mean ± SEM and are analyzed by *t*-test. *p < 0.05, **p < 0.01, vs non-HA group. ns, not significant.

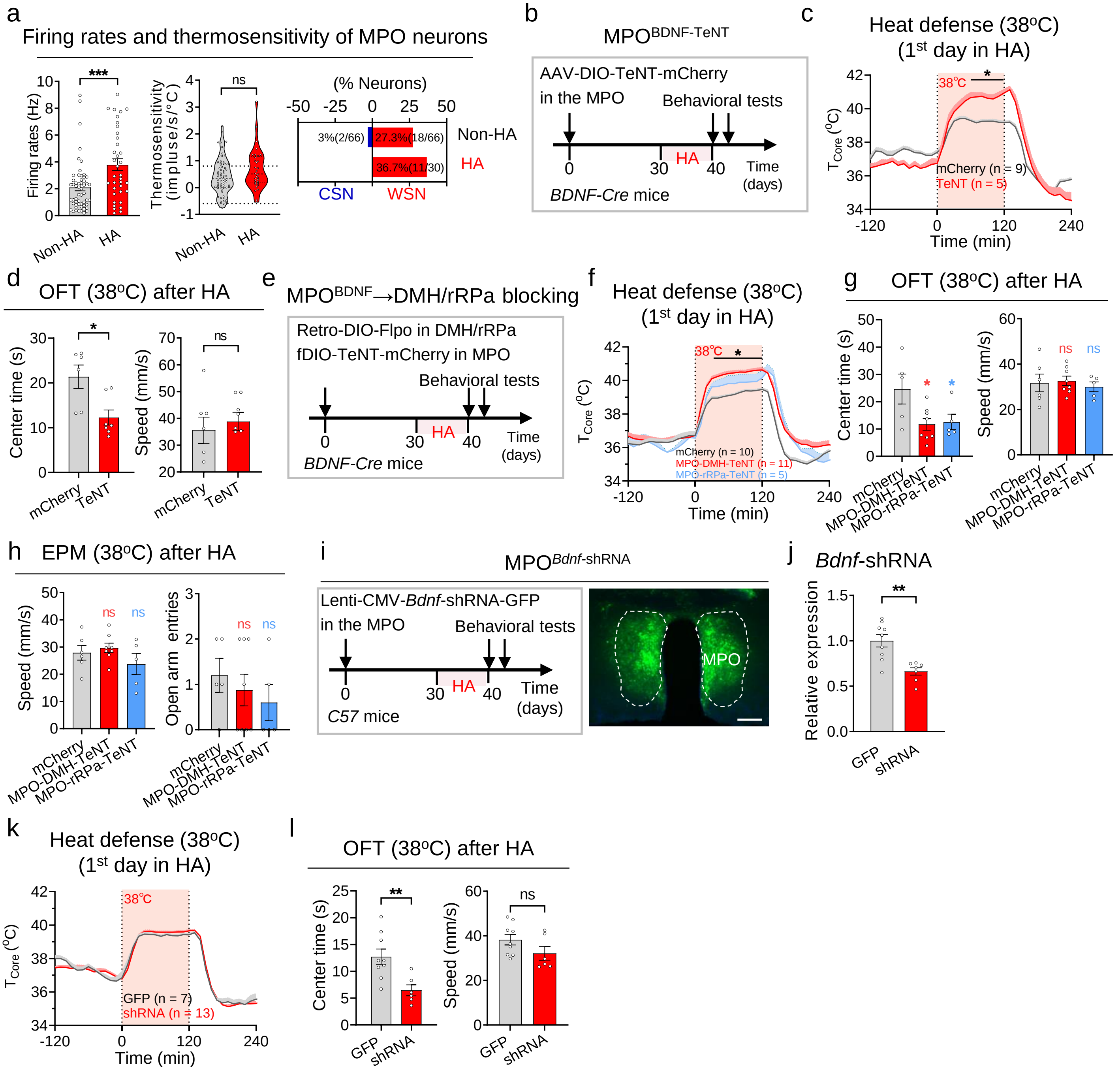

**Fig. S4. BDNF neuron and BDNF molecule in the MPO is required for the HA.**

**a** The basal firing rates at 36ºC, thermosensitivity of neurons in the MPO and the percentage of WSNs and CSNs within all recorded neurons in *C57* mice.

**b** Protocol for HA training after blocking $MPO^{BDNF}$ neurons with TeNT. AAV-DIO-TeNT-mCherry was injected in the MPO of *BDNF-IRES-Cre* mice (termed as $MPO^{BDNF\text{-}TeNT}$, TeNT group). The AAV-DIO-mCherry was used as the control.

**c** $T_{core}$ changes in exposure to 38ºC in the 1st day of HA training in $MPO^{BDNF\text{-}TeNT}$ mice.

**d** OFT after HA training in $MPO^{BDNF\text{-}TeNT}$ mice.

**e** Protocol for HA training after blocking DMH/rRPa-projecting $MPO^{BDNF}$ neurons.

**f** $T_{core}$ changes in exposure to 38ºC in the 1st day of HA training.

**g, h** OFT (**g**) and EPM (**h**) after blocking DMH/rRPa-projecting $MPO^{BDNF}$ neurons.

**i** Design to knockdown *Bdnf* in MPO neurons by injecting Lenti-CMV-*Bdnf*-shRNA into *C57* mice (termed as $MPO^{Bdnf\text{-}shRNA}$, shRNA group) and the representative expression of Lenti-CMV-*Bdnf*-shRNA. The Lenti-CMV-GFP (GFP group) was used as the control.

**j** *Bdnf* mRNA expression in the MPO from $MPO^{Bdnf\text{-}shRNA}$ mice.

**k** $T_{core}$ changes in exposure to 38ºC in the 1st day of HA training in $MPO^{Bdnf\text{-}shRNA}$ mice.

**l** OFT after HA training in $MPO^{Bdnf\text{-}shRNA}$ mice.

All data are shown as mean ± SEM and analyzed by RM ANOVA (**c, f-h, k**) and *t*-test (**a, d, j, l**). *p < 0.05, **p < 0.01 vs corresponding control group. ns, not significant.

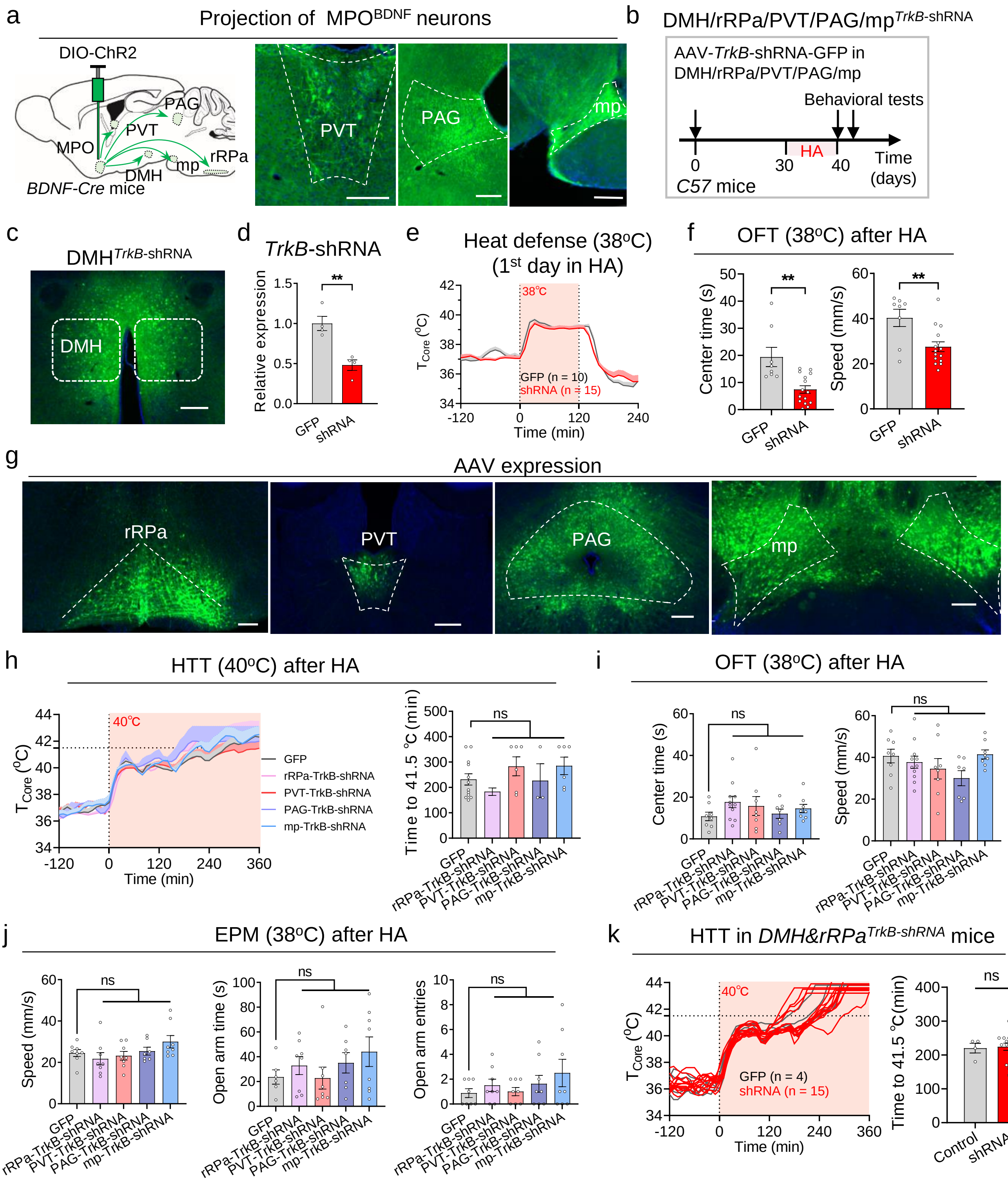


**Fig. S5. Function of TrkB in the HA.**

**a** Axonal expression of MPO$^{BDNF\text{-}ChR2}$ neurons in PVT, PAG and mp. Scale bar, 200 μm.

**b** Protocol for HA training in the *C57* mice after knocking down *TrkB* in downstream regions. AAV-hSyn-Cre & AAV-DIO-shRNA(*TrkB*)-GFP were injected in downstream regions (shRNA group). The mice injected AAV-hSyn-Cre & AAV-DIO-shRNA(*NC*)-GFP (GFP group) were used as the control.

**c** The representative expression of AAV-hSyn-Cre & AAV-DIO-shRNA(*TrkB*)-GFP.

**d** *TrkB* mRNA expression in the DMH from *DMH*$^{TrkB\text{-}shRNA}$ mice.

**e** $T_{core}$ changes in exposure to 38ºC in the 1st day of HA training in *DMH*$^{TrkB\text{-}shRNA}$ mice.

**f** OFT after HA training in *DMH*$^{TrkB\text{-}shRNA}$ mice.

**g** The representative expression of AAV-hSyn-*TrkB*-shRNA. Scale bar, 200 μm.

**h-j** HTT (**h**), OFT (**i**) and EPM (**j**) after HA training.

**k** HTT after HA training in *DMH&rRPa*$^{TrkB\text{-}shRNA}$ mice. AAV-hSyn-Cre & AAV-DIO-shRNA(*TrkB*)-GFP were injected in the DMH & rRPa (termed as *DMH&rRPa*$^{TrkB\text{-}shRNA}$, shRNA group). The mice injected AAV-hSyn-Cre & AAV-DIO-shRNA(*NC*)-GFP (GFP group) were used as the control.

PVT, paraventricular nucleus of the thalamus; PAG, periaqueductal gray; mp, mamillary peduncle. All data are shown as mean ± SEM and analyzed by RM ANOVA(**e, h-j**) and *t*-test (**d, f, k**). $^{**}p < 0.01$ vs corresponding control group. ns, not significant.

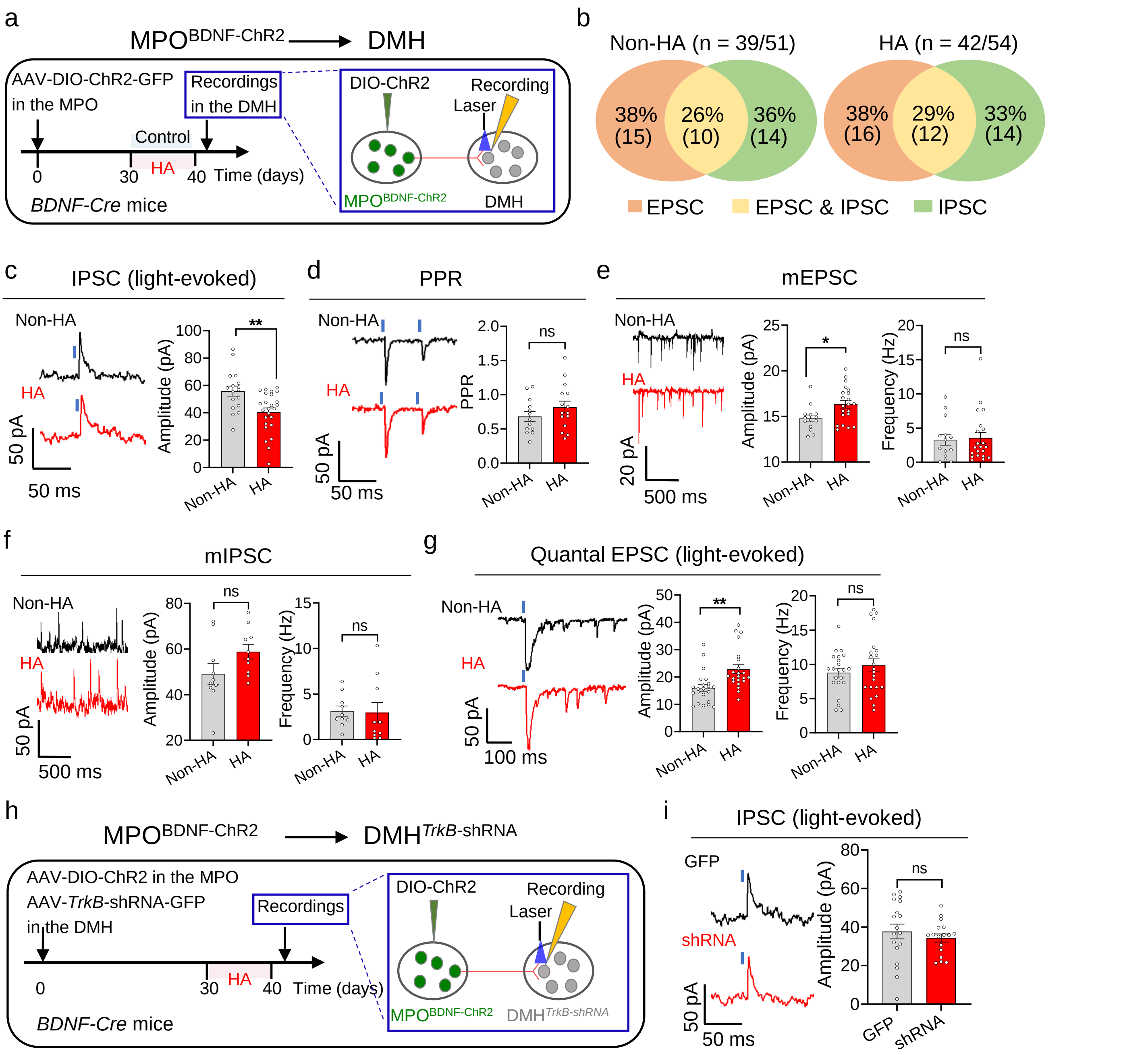


**Fig. S6. HA induces potentiation of the excitatory transmission between MPO$^{BDNF}$ neurons and downstream neurons in the DMH.**

**a** Experimental design to record postsynaptic currents in DMH neurons innervated by MPO$^{BDNF}$ neurons using whole-cell patch configurations. The *MPO$^{BDNF\text{-}ChR2}$* mice without HA training were as the control group.

**b** Numbers of recorded neurons in the DMH following light stimulation of MPO$^{BDNF}$ neurons. 39 neurons out of 51 recorded neurons displayed postsynaptic currents in the control group, while 42 neurons out of 54 recorded neurons displayed postsynaptic currents in the HA group. EPSC/IPSC represents excitatory/inhibitory postsynaptic currents.

**c** Representative traces and amplitudes of IPSCs recorded in DMH neurons following light stimulation (473 nm, 5 ms) of MPO$^{BDNF}$ afferents.

**d** Representative traces of paired pulses (50 ms interval) evoked by light stimulation of MPO$^{BDNF}$ afferents and summary of paired-pulse ratios (PPR).

**e, f** Representative traces of miniature EPSC (mEPSC) (**e**) and miniature IPSC (mIPSC) (**f**) after TTX (1 μM) treatment. The amplitudes and frequency were quantified in the right.

**g** Representative traces and summary of light-evoked quantal EPSCs (le-qEPSCs) recorded in DMH neurons following light stimulation of MPO$^{BDNF}$ afferents in the presence of strontium (5 mM). Quantification of le-qEPSCs was performed on a 300-ms time window and 25-ms after the light pulse.

**h** Experimental design to record postsynaptic currents in DMH neurons innervated by MPO$^{BDNF}$ neurons after knocking down TrkB. AAV-hSyn-Cre & AAV-DIO-shRNA(*TrkB*)-GFP were injected in the DMH of *MPO$^{BDNF\text{-}ChR2}$* mice. The scrambled shRNA group was used as the control.

**i** Representative traces and amplitudes of IPSCs recorded in DMH neurons following light stimulation (473 nm, 5 ms) of MPO$^{BDNF}$ afferents.

All data are shown as mean ± SEM and analyzed by *t*-test. *p < 0.05, **p < 0.01, ***p < 0.001 vs corresponding control group. ns, not significant.
