## Supplementary material for "Heat Acclimation in Mice Requires Preoptic BDNF Neurons and Postsynaptic Potentiation": method

### Supplementary Information List

#### STAR METHODS

- KEY RESOURCES TABLE
- RESOURCE AVAILABILITY
    - Lead contact
    - Materials availability
    - Data and code availability
- EXPERIMENTAL MODEL AND SUBJECT DETAILS
- METHOD DETAILS
    - Stereotaxic brain injection
    - Heat acclimation training
    - Core body temperature measurement
    - Activity and energy expenditure measurement
    - Immunofluorescence staining
    - Heat tolerance test (HTT)
    - Behavioral tests
    - Real-Time PCR
    - *Bdnf* mRNA RNAscope (in-situ hybridization)
    - Light stimulation patterns for optogenetics
    - Brain slice electrophysiology
    - Data quantification and statistical analyses

#### Fig. S1-6

#### REFERENCES FOR SI CITATIONS

### STAR METHODS

#### KEY RESOURCES TABLE

| REAGENT or RESOURCE | SOURCE | IDENTIFIER |
|---|---|---|
| **Antibodies** | | |
| Chicken anti-GFP | Abcam | Cat #ab13970 |
| Rabbit anti-RFP | Abcam | Cat #ab124754 |
| DyLight 488 conjugated goat anti-chicken | Invitrogen | Cat# SA5-10070 |
| Alexa Fluor® 594-AffiniPure Goat Anti-Rabbit IgG (H+L) | Jackson | Cat # 111-585-144 |
| **Chemicals** | | |
| DAPI Fluoromount-G mounting medium | SouthernBiotech | Cat#0100-20 |
| CNQX | Sigma-Aldrich | Cat#C127 |
| AP-5 | Sigma-Aldrich | Cat#A8054 |
| Bicuculline | Alomone | Cat#B-135 |
| TTX | Shanghai Charm-Analysis Technology Co., Ltd | Cat#20-309500 |
| RNAscope® Probe- Mm-Bdnf | acdbio | Cat No. 424821 |
| RNAscope™ 2.5 HD Assay - RED | acdbio | Cat No. 322360 |
| RNA-Protein Co-detection Ancillary Kit | acdbio | Cat No. 323180 |
| **Experimental Models** | | |
| *BDNF-IRES-Cre* | The core facility of Peking Union Medicine College | Reference: [1] |
| *C57BL/6J* | Shanghai Silaike Experiment Animal Co., Ltd. | N/A |
| **Oligonucleotides** | | |
| TrkB shRNA | GATCAATTCAAGAGATTGATC | Reference: [2] |
| Bdnf shRNA | shRNA1:GGTGATGCTCAGCAGTCAAGT;shRNA2:GCAGTATTTCTACGAGACCAA | |

| | | |
|---|---|---|
| **Virus** | | |
| AAV2/9-hEF1a-DIO-mCherry-WPRE-pA | Shanghai Taitool Bioscience Co. | Cat#S0197-9 |
| AAV2/9-hEF1a-DIO-mCherry-P2A-TetTox-WPRE-pA | Shanghai Taitool Bioscience Co. | Cat#S0506-9 |
| AAV2/9-hEF1a-DIO-hChR2(H134R)-EGFP-WPRE-pA | Shanghai Taitool Bioscience Co. | Cat#S0858-9 |
| AAV2/9-hEF1α-DIO-EYFP-WPRE-pA | Shanghai Taitool Bioscience Co. | Cat#S0196-9 |
| AAV2/9-hEF1α-DIO-EGFP-*TrkB*-shRNA | Shanghai Sunbio Medical Biotechnology Co., Ltd. | Cat#BC-1086 |
| rAAV2-Retro-DIO-Flpo-WPRE-hGH polyA | Shanghai Sunbio Medical Biotechnology Co., Ltd. | Cat#PT-0075 |
| AAV2/9-hEF1a-fDIO-mCherry-WPRE-pA | Shanghai Taitool Bioscience Co. | Cat#S0553-9 |
| AAV2/9-hEF1a-fDIO-mCherry-2A-TetTox-WPRE- pA | Shanghai Sunbio Medical Biotechnology Co., Ltd. | Cat#S0551-9-H50 |
| AAV2/8-hSyn-cre-WPRE-pA | Shanghai Taitool Bioscience Co. | Cat#S0278-8-H50 |
| AAV2/9-DIO-EGFP-NC-shRNA | Shanghai Sunbio Medical Biotechnology Co., Ltd. | N/A |
| Lenti-CMV-eGFP-U6-*Bdnf*-shRNA | OBiO Biotechnology Co. | N/A |
| Lenti-CMV-eGFP-NC-shRNA | OBiO Biotechnology Co. | N/A |
| **Coordinates** | | |
| Medial Preoptic Nucleus (MPO) | Anteroposterior AP +0.47 mm, Mediolateral ML ±0.27 mm, Dorsoventral DV -5.17 mm, | *The Mouse Brain in Stereotaxic Coordinates 2nd Edition* |
| dorsomedial hypothalamic nucleus (DMH) | AP -1.20 mm, ML ±0.30 mm, and DV -5.15 mm | *The Mouse Brain in Stereotaxic Coordinates 2nd Edition* |
| rostral raphe pallidus area (rRPa) | AP -6.00 mm, ML 0 mm, | *The Mouse Brain in* |

| | | |
|---|---|---|
| | and DV -6.00 mm | *Stereotaxic Coordinates 2nd Edition* |
| paraventricular nucleus of the thalamus (PVT) | AP -1.00 mm, ML 0 mm, and DV -3.20 mm | *The Mouse Brain in Stereotaxic Coordinates 2nd Edition* |
| periaqueductal gray (PAG) | AP -4.0 mm, ML ±0.75 mm, and DV -3.00 mm | *The Mouse Brain in Stereotaxic Coordinates 2nd Edition* |
| mamillary peduncle (mp) | AP -3.0 mm, ML ±0.70 mm, and DV -5.00 mm | *The Mouse Brain in Stereotaxic Coordinates 2nd Edition* |
| **Software and Algorithms** | | |
| ImageJ bundled with Java 1.8.0_172 | NIH ImageJ | https://imagej.nih.gov/ij/ |
| GraphPad Prism 8 | GraphPad | https://www.graphpad.com/scientific-software/prism/ |
| Clampfit 11.2 | Molecular Devices, LLC | https://www.moleculardevices.com/ |
| Origin 2021 | OriginLab | https://www.originlab.com/ |
| Minianalysis | Synaptosoft | https://www.synaptosoft.com/ |
| MATLAB R2022a | MathWorks | https://www.mathworks.cn/ |
| Polyscan2 | Mightex | https://www.mightexbio.com/ |
| video tracking system (JLBehv-LAM-1) | Shanghai Jiliang Software Technology Co. Ltd | N/A |

#### RESOURCE AVAILABILITY

##### Lead contact

##### Materials availability

This study did not generate new unique reagents or mouse lines.

##### Data and code availability

All the data are available in the main text or the supplementary materials. The data that support the findings of this study were available from the corresponding author upon reasonable request**.** The code that supports the findings of this study was available from the corresponding author upon reasonable request.

#### EXPERIMENTAL MODEL AND SUBJECT DETAILS

##### Experimental mice

Animal care and use conformed to the guidelines of ShanghaiTech University, Shanghai Biomodel Organism Co. Ltd., and Chinese governmental regulations. *BDNF-IRES-Cre* mice were generated through CRISPR/Cas9 genome targeting by the core facility of Peking Union Medicine College, as previously described [1]. *C57BL/6J* mice were purchased from GemPharmatech Co., Ltd. All experiments were performed on adult mice (8-16 weeks old) housed under controlled temperature (22-25 °C, RH 30-40%) in a 12-hour reverse light/dark cycle (dark period: 9 p.m. to 9 a.m.) with ad libitum access to water and chow food (4% fat SPF rodent feed). The animal facility at the National Facility for Protein Science in Shanghai (NFPS) of Zhangjiang Lab and ShanghaiTech University complied with governmental regulations. Unless indicated, adult male mice are used in experiments. All behavioral tests were performed double-blinded during the dark phase. The mice strains were listed in the Key Resources Table.

#### METHOD DETAILS

##### Stereotaxic brain injection

Mice underwent anesthesia with isoflurane, and the target nuclei were located using a stereotaxic frame (David Kopf Instruments, #PF-3983; RWD Life Science, #68030; or Thinker Tech Nanjing Biotech, #SH01A), guided by *The Mouse Brain in Stereotaxic Coordinates 2nd Edition*. To maintain the mice's warmth during surgeries, a feedback heater was utilized. A gradual infusion rate of 20 nl/min delivered 200 nl (unless specified) of AAV (5.0E+12) through a pulled-glass pipette. Post-infusion, the injection needle remained stationary for 10 minutes before being slowly withdrawn. The brain region sites have been listed in the key resources table. For MPO, the optical fiber is inserted 0.3 millimeters above MPO. In order to knockdown *TrkB* in DMH, rRPa, PAG, PVT, and mp, a strategy of co-injecting two viruses was adopted. Specifically, AAV-hSyn-Cre and AAV-DIO-*Trkb*-shRNA-GFP were mixed in a ratio of 1:2, and 200 nL was injected into each of the left and right brain regions.

**Heat acclimation training**

Heat acclimation was conducted using an incubator (Ningbo Jiangnan Instrument Factory, HWS-500) equipped with temperature and relative humidity (RH) control. Three different heat acclimation protocols were employed. In Protocol A, a 14-day uninterrupted training program was implemented. The mice were exposed daily to a temperature of 35°C and a RH of 60 % for a duration of 3 hours. HA Protocol B involved a 20-day sequential rising temperature training program. The temperature gradient ranged from 35°C to 38°C, starting initially at 35°C and increasing by 1°C every 5 days. The RH was maintained at 60 %. The mice were exposed to these heat conditions for 3 hours each day. For HA Protocol C, the mice underwent training for 10 consecutive days during the dark phase. The temperature was set at 38°C, and the RH was maintained at 60 ± 5%. The training sessions lasted for 2 hours per day. Throughout the acclimation period, the mice had ad libitum access to water and food. The mice were housed at temperatures of 22-25°C and a RH of 40 % during other times. The control group was consistently housed under the same conditions of 22-25°C and a RH of 40 %. The mice's weight was systematically recorded at a consistent daily time point. Of note, HA training and testing are highly sensitive to ambient temperature and humidity. These parameters are frequently measured to ensure to the consistency of experimental results.

**Core body temperature measurement**

The core temperature of the mice was continuously monitored using iButton® temperature loggers (DS1922L/T, Analog Devices, USA) at 10-minute intervals, unless stated otherwise. All stimulation and experimental procedures were conducted during the dark phase. The iButton® temperature loggers were implanted in the mice's abdomen at least six days before the start of the experiments. Throughout the duration of the study, the mice were housed in their respective home cages at a temperature of 22-25°C.

**Activity and energy expenditure measurement**

Physical activity, energy expenditure, and body temperature were monitored using the Comprehensive Lab Animal Monitoring System with Temperature Telemetry Transmitter (CLAMS; Columbus Instruments) equipped with G2 E-Mitter transponders. Temperature transponders were surgically implanted in the mice's abdomen 3-5 days prior to the commencement of testing. The mice were given a minimum of 2 days to acclimate to the metabolic chambers before data collection began. Energy expenditure (EE) was recorded at 10-minute intervals, while physical activity was recorded every minute. These measurements were captured and recorded by the CLAMS system, providing detailed information on the mice's metabolic activity, energy expenditure, and body temperature throughout the experimental period.

**Immunofluorescence staining**

Mice were perfused with 50 ml of PBS, followed by 50 ml of 4% PFA. The brain slides were blocked with blocking buffer (2% (v/v) normal goat serum, 2.5% (w/v) bovine serum albumin, 0.1%

(v/v) Triton X-100, and 0.1% (w/v) NaN3 in PBS) for 1 hour at room temperature, and then incubated with primary antibodies (chicken-anti-GFP or rabbit anti-RFP, 1:1000) for 1 day at 4°C. Subsequently, the samples were washed three times with PBST (PBS with 0.1% Triton X-100) before being incubated with secondary antibodies (488-goat anti-chicken or 594-goat anti-rabbit, 1:1,000) overnight at 4°C. After washing with PBST three times, the slide was sealed.

**Heat tolerance test (HTT)**

Animals were transferred from room temperature to the 40°C, 40%RH chamber to participate in the heat tolerance test which lasted up to 10 hours. The mice had free access to food and water, and their body temperature was continuously monitored. The cut-off threshold for body temperature was 41.5°C [3,4]. Some mice died after prolonged exposure to 40°C, and their $T_{core}$ after death were replaced with the highest $T_{core}$ to produce a flattened trace shown on figures.

**Behavioral tests**

**Open field test (OFT).** To assess anxiety-like behavior, mice were tested in the open field test (OFT) according to a previously described protocol [5,6]. The mice were placed in a 50×50×50 cm top-opened box under dim light conditions (approximately 10-20 lux) at a thermal neutral temperature (25°C). In the control (non-acclimation) group, mice were placed in the center of the arena and allowed to freely explore for 5 minutes. For the acclimated group, mice were stimulated with 2-hour heat exposure (38°C, 60% RH) prior to the OFT test. The behavior of the animals was recorded and subsequently analyzed using a video tracking system (JLBehv-LAM-1, Shanghai Jiliang Software Technology Co. Ltd).

**Elevated plus maze (EPM).** The elevated plus maze (EPM) consisted of a plus-shaped apparatus with a central platform (5×5 cm), two open arms (25×5 cm), two closed arms (25×5×5 cm), and elevated 50 cm from the floor, illuminated with dim light (approximately 10-20 lux)[7]. The testing was conducted in a blind fashion, where each mouse was placed on the central platform facing the open arm. The following standards in the EPM behavior analysis software (JLBehv-LAM-1, Shanghai Jiliang Software Technology Co. Ltd) had been set: the software determines the position of the mouse by using the Geometric Centroid of its body, identified from the highlighted areas in the image (excluding the tail). Regarding the standard for "entry", generally, more than 80% of the mouse's body enters the open arm[8-10]. According to the body length of adult mice and the length of the open arm of the elevated plus maze, this distance in the software is set at 50 mm[11-13]. And only when this positioning point fully enters the open arm by 50 mm will the software record an entry. The movement of the mice was recorded and analyzed for 5 minutes using a video tracking system.

**Hot plate test.** The hot plate test was utilized to measure the heat nociception threshold in mice. The hot plate was maintained at 55 ± 0.5°C, and the mice were placed unrestrained on the plate until a nociceptive response was observed. Nociceptive responses include paw withdrawal or licking, stamping, leaning, or jumping. The mouse was removed from the hot plate, even if none of the above-mentioned behaviors were shown within 30 seconds, to prevent any potential skin scalding.

**Y maze.** The Y-maze device comprises three adjacent arms set at an angle of 120°, forming a central equilateral triangle. Each arm is labeled with a distinct shape (square, circle, or triangle) for differentiation. The device is situated in a controlled environment within a constant temperature and humidity incubator to ensure stability during the experiment. During the pre-adaptation phase on the first day, one arm is obstructed by a movable partition, and mice are placed in the initial arm for a 15-minute adaptation period. On the second testing day, the movable partition is removed, and mice are still introduced from the initial arm. Video recording begins upon placement, and after 5 minutes, it is stopped, followed by gentle removal of the mice. Thorough cleaning of the entire Y-maze device with distilled water and 75% alcohol is performed after each mouse's test, and subsequent testing is conducted once the incubator's temperature and humidity have stabilized.

**Temperature preference test.** The temperature selection box consists of two separate chambers with metal temperature control panels at the bottom. To assess the ambient temperature preference of the mice, the temperature selection box was placed in a dedicated behavioral sound insulation box, and the activities of the mice were monitored and recorded using infrared cameras. Prior to the formal experiment, the mice underwent 2 hours of daily environmental adaptation training for 2 days. During the training, both chambers were maintained at room temperature (24 ± 1℃). For the formal test, after stabilizing the temperature of the bottom plate, the temperature selection box was introduced in the same order and at the same time as during the adaptation training. A 30-minute video recording was conducted during the test. Following the test, the device was thoroughly cleaned using water and alcohol. The residence time of the mice in the two chambers was analyzed to determine their preference rate.

**Thermoregulatory behavior recording.** Mice were individually housed in standard cages for heat acclimation prior to testing. Subsequently, each mouse was subjected to a thermal stimulus of 40°C for 20 minutes while being video recorded. For "extension", when a mouse extends limbs and remains still for more than 2 seconds, it's considered extension. Record the start time and total time. For "grooming", record the start time and total time.

**Real-Time PCR**

Real-time PCR (qPCR) was performed to quantify gene expression levels in the brain tissue, specifically the POA region. Total RNA was isolated using Trizol (Sangon, #B511311-0100), followed by DNase I treatment (Takara, #RR047A). Reverse transcription of 1.0 µg of total RNA was carried out using PrimeScriptÔ RT Master Mix (Takara, #RR047A). qPCR was conducted in triplicate with an SYBR Green PCR kit (Takara, #RR820A) in a 10 µl reaction volume. Mouse GAPDH was utilized as the endogenous control for normalization of sample values, and expression levels were calculated using the 2^(-ΔΔCq) method. The primer sequences used in this study were as follows:

*Bdnf* (TCATACTTCGGTTGCATGAAGG, AGACCTCTCGAACCTGCCC), *TrkB* (CTGGGGCTTATGCCTGCTG, AGGCTCAGTACACCAAATCCTA), *Ucp1* (GGCCCTTGTAAACAACAAAATAC, GGCAACAAGAGCTGACAGTAAAT).

**_Bdnf_ mRNA RNAscope (_in-situ_ hybridization)**

To detect BDNF expression, mRNA in situ hybridization using the RNAscope 2.5 HD-Red Assay (ACD, Inc., Newark, NJ, USA) with probes specific to murine Mm-Bdnf (Cat No. 424821) was performed on PFA-fixed (10%) brain sections. Mice were deeply anesthetized, followed by transcardial perfusion with 4% paraformaldehyde. The brains were dissected, and then dehydrated in 20% and 30% sucrose for one day at 4°C sequentially. Cryosection was performed and sections (16 μm) were placed onto SuperFrost Plus slides, dried at 20°C for 1 hour, and stored at -80°C. Before probe incubation, the sections were pretreated with hydrogen peroxide for 10 minutes at room temperature, followed by Protease treatment for 30 minutes at 40°C. Washing with 1× PBS for 5 minutes to remove OCT. Hydrogen peroxide was applied to each section and incubated at room temperature for 10 minutes. Subsequently, boiling 1× target repair reagent was used for 5 minutes to repair the sections, followed by washing with distilled water and 100% ethanol. Protease was then applied to the hydrophobic circle on each brain slice and incubated at 40°C for 30 minutes, followed by washing with distilled water. After the pretreatment steps, the brain sections were incubated with the individual probes for 2 hours, and the standard RNAscope protocol was followed, including sequential incubations with Amp 1 (at 40°C for 30 minutes), Amp 2 (at 40°C for 15 minutes), Amp 3 (at 40°C for 30 minutes), Amp 4 (at 40°C for 15 minutes), Amp 5 (at room temperature for 30 minutes), and Amp 6 (at room temperature for 15 minutes). Finally, the RED working solution was added to detect the signal. Following rinsing, the slides were cover-slipped with DAPI Fluoromount-G mounting medium (SouthernBiotech, #0100-20). Image acquisition was performed using a Nikon CSU Sora 2 Camera microscope with a 20x objective (Nikon, CSU-

W1) or an Olympus VS120 Virtual Microscopy Slide Scanning System (Olympus, VS120-S6-W). Cell numbers were quantified using ImageJ software (NIH ImageJ).

**Light stimulation patterns for optogenetics**

Light stimulation patterns for optical stimulation were implemented using a 473-nanometer laser (Inper, China). The light intensity emitted from the fiber tip was measured using a power meter (PM20A, Thorlabs). In the case of ChR2-expressing mice, light pulses of 10-ms duration and 4 mW power were administered via a fiber optic probe at a frequency of 1 Hz. The light stimulation protocol was initiated three days before the heat endurance phase.

**Brain Slice Electrophysiology**

Brain slice electrophysiology recording was performed on male *C57BL/6* mice and *BDNF-IRES-Cre* mice. Mice were anesthetized with tribromoethanol (100 mg/kg, i.p.) and perfused transcardially with ice-cold oxygenated (95% $O_2$/5% $CO_2$) NMDG ACSF solution, which included 93 mM NMDG, 93 mM HCl, 2.5 mM KCl, 1.25 mM $NaH_2PO_4$, 10 mM $MgSO_4 \cdot 7H_2O$, 30 mM $NaHCO_3$, 25 mM glucose, 20 mM HEPES, 5 mM sodium ascorbate, 3 mM sodium pyruvate, and 2 mM thiourea. After perfusion, the brain was rapidly dissected out and immediately transferred into an ice-cold oxygenated NMDG ACSF solution. Then, we sectioned the coronal plane of brain tissue at 300 μm in the same buffer using a vibratome (VT1200S, Leica). The brain slices containing the POA were incubated in oxygenated NMDG ACSF at 32°C for 10-15 min and then transferred to a normal oxygenated solution of ACSF (126 mM NaCl, 2.5 mM KCl, 1.25 mM $NaH_2PO_4$, 2 mM $MgSO_4 \cdot 7H_2O$, 10 mM glucose, 26 mM $NaHCO_3$, 2 mM $CaCl_2$) at room temperature for 1 hour. All chemicals used in slice preparation were purchased from Sigma-Aldrich (St. Louis, MO, USA). Slices were transferred to the recording chamber that was submerged and superfused with ACSF at a rate of 3 mL/min at the neutral temperature (35-37°C).

Whole-cell patch-clamp recordings were made from neurons visualized with an Olympus BX61W1 microscope (equipped with GFP and mCherry filters) using infrared video microscopy and differential interference contrast optics. BDNF neurons were identified by GFP fluorescence. The recording pipettes (4-8 MΩ) were prepared with a micropipette puller (P2000, Sutter Instrument; USA). For whole-cell recording, the pipettes were filled with ACSF solution containing 133 mM potassium gluconate, 18 mM NaCl, 0.6 mM EGTA, 10 mM HEPES, 2 mM Mg·ATP, and 0.3 mM Na3·GTP (pH:7.2, 280 mOsm).

The thermoelectric assembly allowed the tissue slices' temperature to be periodically varied 3-5°C above and below the thermoneutral temperature (35-37°C) to characterize the thermosensitivity of each recorded neuron. The firing activity of neurons was continuously recorded during cyclic temperature changes. To test the intrinsic neural thermosensitivity, synaptic transmission blockers of AMPA-receptors (CNQX, 10 μM), NMDA-receptors (AP-5, 50 μM), and GABAA-receptors (bicuculline, 25 μM) were added to the chamber throughout the entire recording. Thermosensitivity (impulses/s/°C) was defined by the slope of the linear regression line (thermal coefficient) plotted by the firing rate as a function of temperature. This plot was determined over a temperature range of at least 3°C. Based on the thermal coefficient, neurons were classified as warm-sensitive neurons (WSNs, thermal coefficient ≥ 0.8 impulses/s/ ºC), cold-sensitive neurons (CSNs, thermal coefficient ≤ -0.6 impulses/s/ºC), and temperature-insensitive neurons (TINs, thermal coefficient < 0.8 but > -0.6 impulses/s/ ºC) [14-16]. Considering the importance of intrinsic thermosensitivity in heat defense and its proposed role in HA, we first recorded the basal firing rates at 36ºC.

To photostimulate ChR2-expressing fibers, an LED light source (473 nm) was focused onto the back aperture of the microscope objective (Mightex polygon400), producing a beam of light that targets a single cell. The light output was controlled by a programmable pulse stimulator (Mightex-Polyscan2) and pClamp 11.2 software (Molecular Devices). For light evoked-qEPSC (le-qEPSCs) recordings, the membrane potential was clamped at $V_h$ = -70 mV and the protocol consisted of one blue light pulse (473-nm wavelength, 5 ms) applied every 20 seconds. le-qEPSCs measurements were performed in ACSF containing 126 mM NaCl, 2.6 mM KCl, 1.2 mM $NaH_2PO_4$, 10 mM glucose, 21.4 mM $NaHCO_3$, 1.2 mM $MgCl_2$ $6H_2O$, 5mM $SrCl_2$. A 300-ms time window 25 ms after the light pulse was used for analysis of the evoked quantal events [17]. Results from 5 sweeps were averaged for analysis. For le-EPSC and le-IPSC recordings, the membrane potential was clamped at $V_h$ = -70 mV or 40 mV, respectively. The detection protocol consisted of blue light pulses (473-nm wavelength, 5 ms). For paired-pulse ratio (PPR), two light pulses were administered 50 ms apart every 20 seconds. For analysis, 3-5 sweeps were averaged.

For miniature excitatory postsynaptic current (mEPSC) and miniature inhibitory postsynaptic current (mIPSC), the membrane potential was clamped at $V_h$ = -70 mV or 40 mV, respectively. All experiments were performed in the presence of tetrodotoxin (TTX, 1 μM). All recordings were acquired using a Multiclamp 700B amplifier, and signals were low-pass filtered at 3 kHz and digitized at 10 kHz (DigiData 1550B, Molecular Devices). The physiological data were analyzed using Clampfit 11.2 software (Molecular Devices) and Mini Analysis Program (Synaptosoft).

**Data quantification and statistical analyses**

Data collection and analyses were conducted in a blinded manner to the experimental conditions. Statistical analyses of the experimental data were performed using *t*-test and analysis of variance (ANOVA), as indicated. The "n" value used in these analyses represents the number of mice or cells included in the study.

#### References

1. Luo, F., Mu, Y., Gao, C., Xiao, Y., Zhou, Q., Yang, Y., Ni, X., Shen, W.L., and Yang, J. (2019). Whole-brain patterns of the presynaptic inputs and axonal projections of BDNF neurons in the paraventricular nucleus. J Genet Genomics 46, 31-40. 10.1016/j.jgg.2018.11.004.
2. Bartkowska, K., Paquin, A., Gauthier, A.S., Kaplan, D.R., and Miller, F.D. (2007). Trk signaling regulates neural precursor cell proliferation and differentiation during cortical development. Development 134, 4369-4380. 10.1242/dev.008227.

3. Wilkinson, D.A., Burholt, D.R., and Shrivastava, P.N. (1988). Hypothermia following whole-body heating of mice: effect of heating time and temperature. Int J Hyperthermia 4, 171-182. 10.3109/02656738809029307.
4. Koh, Y.H. (2018). Heat Stroke with Status Epilepticus Secondary to Posterior Reversible Encephalopathy Syndrome (PRES). Case Rep Crit Care 2018, 3597474. 10.1155/2018/3597474.
5. Li, C., Yan, Y., Cheng, J., Xiao, G., Gu, J., Zhang, L., Yuan, S., Wang, J., Shen, Y., and Zhou, Y.D. (2016). Toll-Like Receptor 4 Deficiency Causes Reduced Exploratory Behavior in Mice Under Approach-Avoidance Conflict. Neurosci Bull 32, 127-136. 10.1007/s12264-016-0015-z.
6. Zhao, Z.D., Chen, Z., Xiang, X., Hu, M., Xie, H., Jia, X., Cai, F., Cui, Y., Chen, Z., Qian, L., et al. (2019). Zona incerta GABAergic neurons integrate prey-related sensory signals and induce an appetitive drive to promote hunting. Nature neuroscience 22, 921-932. 10.1038/s41593-019-0404-5.
7. Pi, G., Gao, D., Wu, D., Wang, Y., Lei, H., Zeng, W., Gao, Y., Yu, H., Xiong, R., Jiang, T., et al. (2020). Posterior basolateral amygdala to ventral hippocampal CA1 drives approach behaviour to exert an anxiolytic effect. Nat Commun 11, 183. 10.1038/s41467-019-13919-3.
8. Peres, D.S., Viero, F.T., Rodrigues, P., de Barros Bernardes, L., da Silva, N.A.R., Lima, I.R., Martins, G., Silveira, P.C.L., de Amorim Ferreira, M., Silva, A.M., et al. (2023). Characterization of Depression- and Anxiety-Like Behaviours in a Mouse Model of Relapsing-Remitting Multiple Sclerosis. J Neuroimmune Pharmacol 18, 235-247. 10.1007/s11481-023-10080-z.
9. Rabbani, M., Sajjadi, S.E., and Jalali, A. (2005). Hydroalcohol extract and fractions of Stachys lavandulifolia Vahl: effects on spontaneous motor activity and elevated plus-maze behaviour. Phytother Res 19, 854-858. 10.1002/ptr.1701.
10. Saitoh, A., Nagayama, Y., Yamada, D., Makino, K., Yoshioka, T., Yamanaka, N., Nakatani, M., Takahashi, Y., Yamazaki, M., Shigemoto, C., et al. (2022). Disulfiram Produces Potent Anxiolytic-Like Effects Without Benzodiazepine Anxiolytics-Related Adverse Effects in Mice. Front Pharmacol 13, 826783. 10.3389/fphar.2022.826783.
11. Hong, Y., Hu, J., Zhang, S., Liu, J., Yan, F., Yang, H., and Hu, H. (2024). Integrative analysis identifies region- and sex-specific gene networks and Mef2c as a mediator of anxiety-like behavior. Cell Rep 43, 114455. 10.1016/j.celrep.2024.114455.
12. Yang, H., Xie, J., Mu, W., Ruan, X., Zhang, J., Yao, L., Diao, Z., Wu, M., Li, Y., Ren, W., and Han, J. (2020). Tea polyphenols protect learning and memory in sleep-deprived mice by promoting AMPA receptor internalization. Neuroreport 31, 857-864. 10.1097/wnr.0000000000001462.
13. Zhang, K., Li, Y.J., Guo, Y., Zheng, K.Y., Yang, Q., Yang, L., Wang, X.S., Song, Q., Chen, T., Zhuo, M., and Zhao, M.G. (2017). Elevated progranulin contributes to synaptic and learning deficit due to loss of fragile X mental retardation protein. Brain 140, 3215-3232. 10.1093/brain/awx265.
14. Ranels, H.J., and Griffin, J.D. (2003). The effects of prostaglandin E2 on the firing rate activity of thermosensitive and temperature insensitive neurons in the ventromedial preoptic area of the rat hypothalamus. Brain Res 964, 42-50. 10.1016/s0006-8993(02)04063-5.
15. Griffin, J.D., and Boulant, J.A. (1995). Temperature effects on membrane potential and input resistance in rat hypothalamic neurones. J Physiol 488 ( Pt 2), 407-418. 10.1113/jphysiol.1995.sp020975.
16. Zhou, Q., Fu, X., Xu, J., Dong, S., Liu, C., Cheng, D., Gao, C., Huang, M., Liu, Z., Ni, X., et al. (2023). Hypothalamic warm-sensitive neurons require TRPC4 channel for detecting

internal warmth and regulating body temperature in mice. Neuron 111, 387-404.e388. 10.1016/j.neuron.2022.11.008.

17. Grzelka, K., Wilhelms, H., Dodt, S., Dreisow, M.L., Madara, J.C., Walker, S.J., Wu, C., Wang, D., Lowell, B.B., and Fenselau, H. (2023). A synaptic amplifier of hunger for regaining body weight in the hypothalamus. Cell metabolism. 10.1016/j.cmet.2023.03.002.
